# BioCCP.jl: Collecting Coupons in combinatorial biotechnology

**DOI:** 10.1101/2021.07.09.451763

**Authors:** Kirsten Van Huffel, Michiel Stock, Bernard De Baets

## Abstract

In combinatorial biotechnology, it is crucial for screening experiments to sufficiently cover the design space. In the `BioCCP.jl` package, we provide functions for minimum sample size determination based on the mathematical framework coined the Coupon Collector Problem. `BioCCP.jl`, including source code, documentation and a Pluto notebook is available at https://github.com/kirstvh/BioCCP.

## Combinatorial biotechnology and the Coupon Collector Problem

Biological entities, such as proteins, genetic circuits and genomes, follow the modularity principle, as they can be decomposed into basic functional parts or modules. Synthetic biologists exploit this organizational feature to engineer biological systems that can perform new, specific tasks (1). Oftentimes, a combinatorial approach is adopted, where diverse libraries of genetic variants are generated by assembling and recombining modules (2). To this end, recent advances in recombinant DNA techniques have revolutionized the ability to generate combinatorial constructs in a high-throughput manner (3). Integration of these constructs into host cells brings about novel phenotypes with rich biological activities, of which the optimal variants are selected using screening techniques. For instance, a biotechnologist might design a library of enzyme variants in which the protein domains are diversified and reorganized to enable screening for the set of domains that optimizes enzyme activity (4). Alternatively, in the case of combinatorial CRISPR experiments in plants, combinatorial constructs of guideRNAs are introduced in cells to create a large number of plants with different combinations of targeted knockouts in the genome, to then select plants with the desired phenotype (5).

With an increasing number of available modules to compose biological designs, a combinatorial explosion of possibilities arises. Hence, usually only a limited subset of a combinatorial library can be screened in the laboratory. A crucial decision here is to determine a sample size that guarantees a sufficient coverage of the design space. For example, in the case of the combinatorial engineering of a protein by randomizing protein domains, one aims to determine how many proteins from the library must be sampled in order to collect the complete set of protein domains. For combinatorial CRISPR knockouts in plants, a researcher wants to gain insight into the minimum required number of plants to realize each possible gene knockout at least once. In general, before designing the experiments, the researcher is interested to know the expected number of designs that needs to be assembled to observe every module at least once. This conundrum is addressed by the Coupon Collector Problem (CCP).

In probability and statistics, the CCP describes a situation where there are *n* different types of “coupons” of which a collector tries to obtain a complete set. For this purpose, the collector has to repeatedly sample one coupon at a time from a population (e.g., from cereal boxes) with replacement. A mathematical question that naturally arises from this experiment is the expected number of draws that are required to complete the collection. Importantly, if every coupon is equiprobable, one has to draw 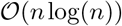 samples. This standard formulation of the CCP and several variants thereof have been subjected to extensive research by scientists over the past decades (6–11) and are still subject of ongoing study in practical settings, such as computational time analysis of algorithms (12), ecological studies (13, 14) and linguistics (15). However, the practicality of the CCP in the field of combinatorial biotechnology has not yet been thoroughly highlighted. In this application note, we clarify the importance of the CCP for experimental design in biotechnology, in particular its potential to define a theoretical lower bound for the sample size in screening experiments. To facilitate practical application by biotechnologists, we provide the `BioCCP.jl` package in Julia and a corresponding interactive Pluto notebook (https://github.com/kirstvh/BioCCP). Before rephrasing minimum sample size determination in combinatorial biotechnology as a CCP, two peculiarities regarding biotechnological designs should be emphasized. Firstly, a design can be an assembly of several modules (e.g., a protein consisting of multiple protein domains). Hence, sampling one design can correspond to drawing multiple modules instead of just one module. Secondly, the modules might not occur with equal frequency (e.g., heterogeneous concentrations of DNA fragments during library generation, deliberately or accidentally due to inaccurate lab techniques). Therefore, finding a minimum sample size differs from the standard formulation of the CCP with equiprobable coupons, and translates to a CCP with unequal coupon probabilities (9, 11).

## Expected minimum number of designs

In the following, we exploit the CCP with unequal coupon probabilities to compute the expected minimum sample size to collect designs from a combinatorial library. Consider the designs in this library to be created by randomly combining *r* modules (with replacement) from a collection of *n* modules (Fig. 1A). Hence, a design is assumed to be a multi-set of *r* modules. The modules exhibit (unequal) probabilities (*p*_1_, …, *p*_n_) with 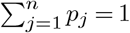 during library generation. The goal is to randomly collect a number of designs so that all *n* modules are observed at least once in the entire collection. A general formula for the expected minimum number of designs can be derived by considering the random incorporation of modules into combinatorial constructs as independent Poisson processes. We refer to the work of (11) for a description on Poissonization of the CCP. Based on their findings, we can write the formula for the expected minimum number of designs as:

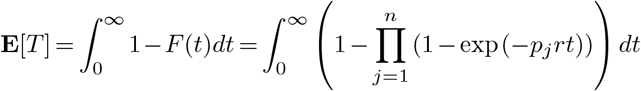

with *n* the number of modules, *p*_j_ the probability of module *j*, *r* the number of modules per design and *t* the number of designs. *T* is a variable representing the minimum number of designs to encounter all possible modules at least once for a specific experiment. *F* (*t*) denotes the cumulative distribution of *T*, *i.e.*, the probability that the required number of designs is smaller than or equal to *t*, and will be further addressed as the “success probability” (16). Fig.1B illustrates that **E**[*T*] can be obtained by integrating 1 − *F* (*t*) over the entire domain. The latter also allows for the calculation of the variance of *T* by computing the second moment (11).

**Fig. 1.**
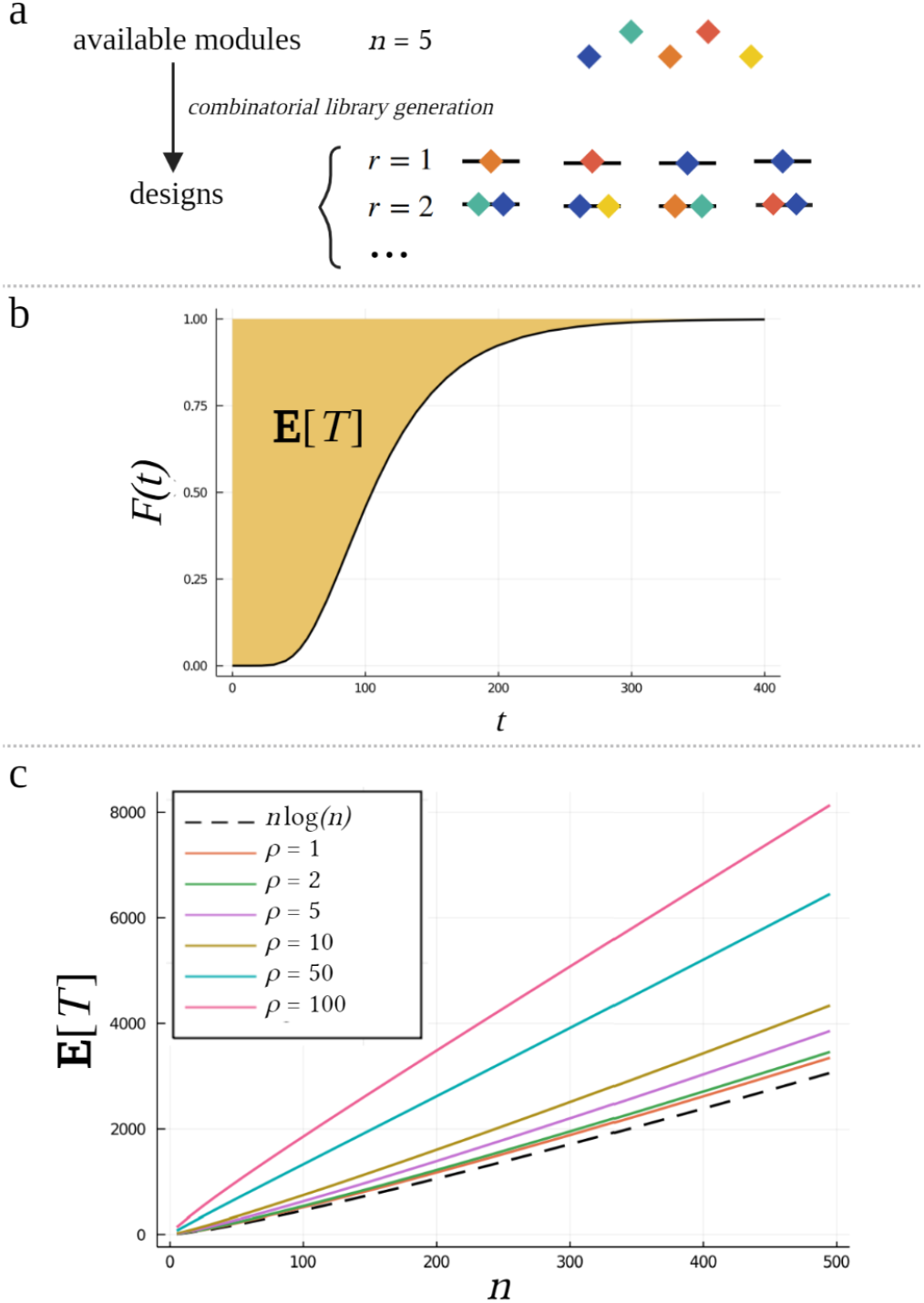
(a) Designs in combinatorial biotechnology; (b) Calculation of the expected minimum number of designs by integrating 1 *− F* (*t*); (c) The expected minimum number of designs in function of the number of available modules for different ranges of module probabilities.

The probabilities of the biological modules (*p*_1_,…, *p*_n_) are often not known exactly. In such case, we propose to define the range of probabilities through the ratio *ρ* = max*_j_ p_j_* / min*_j_ p_j_*, and fix the probability vector through a power law series (according to Zipf’s law):

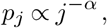

with *j* the rank of the module when the probabilities (*p*_1_,…, *p*_n_) are sorted from high to low and *α* a constant determined by *ρ* and *n*. This power law renders the probability of a module *p_j_* to be inversely proportional to its index *j*. In Fig. 1C, the expected minimum number of designs in function of the total number of modules in the design space is visualized for different values of *ρ*. It is clear that the curve of the expected minimum number of designs moves upward as this ratio increases.

## The BioCCP.jl package

With the `BioCCP.jl` package and its accompanying Pluto notebook, we intend to increase understanding about minimum sample sizes for combinatorial screening experiments by providing the following functionality. After defining the specifics of a combinatorial design setting of interest (the number of modules in the design space, the number of modules per design and the module probabilities), `BioCCP.jl` enables: (i) calculating the minimum expected number of designs to observe all modules at least *m* times and the corresponding standard deviation (see the generalization of the CCP for collecting multiple complete sets described by Doumas and Papanicolaou (11) and Boneh and Hofri (16)); (ii) computing the success probability to encounter all modules at least *m* times for a fixed sample size; ( iii) examining the expected observed fraction of the total number of available modules in function of sample size; (iv) studying the influence of different module probability distributions on the aforementioned statistics, and; (v) calculating the probability that a specific module occurs a given number of times in a fixed sample size (16). Both working with user-specific probabilities of biological modules as well as prototypical probability distributions (e.g., Zipf’s law) is enabled. We anticipate that unlocking this mathematical framework to biotechnologists will assist in obtaining a deeper understanding of statistical properties of biological datasets, facilitating sound experimental design and proper decision making from screening experiments.

## Bibliography

1. Ernesto Andrianantoandro, Subhayu Basu, David K Karig, and Ron Weiss. Synthetic biology: new engineering rules for an emerging discipline. Molecular Systems Biology, 2(1): 2006.0028, 2006.

2. Michael J Smanski, Swapnil Bhatia, Dehua Zhao, YongJin Park, Lauren BA Woodruff, Georgia Giannoukos, Dawn Ciulla, Michele Busby, Johnathan Calderon, Robert Nicol, et al. Functional optimization of gene clusters by combinatorial design and assembly. Nature Biotechnology, 32(12):1241–1249, 2014.

3. Ran Chao, Yongbo Yuan, and Huimin Zhao. Recent advances in DNA assembly technologies. FEMS Yeast Research, 15(1):1–9, 2015.

4. Hans Gerstmans, Dennis Grimon, D Gutiérrez, Cédric Lood, Ana Rodríguez, Vera van Noort, Jeroen Lammertyn, Rob Lavigne, and Yves Briers. A VersaTile-driven platform for rapid hit-to-lead development of engineered lysins. Science Advances, 6(23):eaaz1136, 2020.

5. Norbert Bollier, Rafael Andrade Buono, Thomas B Jacobs, and Moritz K Nowack. Efficient simultaneous mutagenesis of multiple genes in specific plant tissues by multiplex CRISPR. Plant Biotechnology Journal, 19(4):651–653, 2021.

6. Donald J Newman. The double dixie cup problem. The American Mathematical Monthly, 67(1):58–61, 1960.

7. Paul Erdös and Alfréd Rényi. On a classical problem of probability theory. Magyar Tud. Akad. Mat. Kutato Int. Kozl, 6:215–219, 1961.

8. William Feller. An Introduction to Probability Theory and its Applications, volume II. John Wiley & Sons, 1971.

9. Lars Holst. On Birthday, Collectors’, Occupancy and Other Classical Urn Problems. International Statistical Review/Revue Internationale de Statistique, 54(1):15–27, 1986.

10. Philippe Flajolet, Daniele Gardy, and Loÿs Thimonier. Birthday paradox, coupon collectors, caching algorithms and self-organizing search. Discrete Applied Mathematics, 39(3):207–229, 1992.

11. Aristides V Doumas and Vassilis G Papanicolaou. The coupon collector’s problem revisited: generalizing the double dixie cup problem of Newman and Shepp. ESAIM: Probability and Statistics, 20:367–399, 2016.

12. Yu-an Zhang, Xiaofeng Qin, Qinglian Ma, Minghao Zhao, Satoru Hiwa, Tomoyuki Hiroyasu, and Hiroshi Furutani. Markov chain analysis of evolutionary algorithms on OneMax function–From coupon collector’s problem to (1+ 1) EA. Theoretical Computer Science, 820:26–44, 2020.

13. Noemi Zoroa, Emmanuel Lesigne, María-José Fernández-Sáez, Procopio Zoroa, and Jérôme Casas. The coupon collector urn model with unequal probabilities in ecology and evolution. Journal of The Royal Society Interface, 14(127):20160643, 2017.

14. Arik Kershenbaum, Todd M Freeberg, and David E Gammon. Estimating vocal repertoire size is like collecting coupons: a theoretical framework with heterogeneity in signal abundance. Journal of Theoretical Biology, 373:1–11, 2015.

15. Lukun Zheng and Huiqiang Zheng. Authorship Attribution via Coupon-Collector-Type Indices. Journal of Quantitative Linguistics, 27(4):321–333, 2020.

16. Arnon Boneh and Micha Hofri. The coupon-collector problem revisited–a survey of engineering problems and computational methods. Stochastic Models, 13(1):39–66, 1997.

